# Death receptor 6 does not regulate axon degeneration and Schwann cell injury responses during Wallerian degeneration

**DOI:** 10.1101/2025.07.21.665928

**Authors:** Bogdan Beirowski, Haoran Huang, Elisabetta Babetto

## Abstract

Axon degeneration (AxD), accompanied by glial remodeling, is a pathological hallmark of many neurodegenerative diseases, leading to the disruption of neuronal connectivity [1–3]. Understanding the mechanisms in neurons and glia that regulate AxD is essential for developing therapeutic strategies to prevent or mitigate axon loss. Wallerian degeneration (WD) is a well-established model to study the mechanisms of nerve injury-induced AxD, glial responses, and axon-glia interactions. We recently showed that Schwann cells (SCs), the axon-associated glia of the peripheral nervous system, exert protective effects on axons through their rapid metabolic injury response [4]. Enhancing this SC response promotes axon protection during WD [4]. A prior study reported that eliminating the orphan tumor necrosis factor receptor DR6 (death receptor 6, Tnfrsf21) strongly delays AxD and alters SC injury responses during WD [5], suggesting a possible intersection with our findings. Here, we rigorously revisit the role of DR6 in WD using two independent DR6 knockout mouse lines including the same model used in the previous study. Surprisingly, in striking contrast to the earlier report, we observed no impact of DR6 deletion on AxD kinetics or SC injury responses across a range of WD assays. Moreover, injured axons in primary neuronal cultures lacking DR6 degenerated at a similar rate as wild-type axons. We conclude that DR6 is dispensable for the regulation of AxD and glial nerve injury responses during WD. Our data argue that any therapeutic benefit from DR6 suppression in neurodegeneration models occurs through mechanisms independent of WD.

## Results

### AxD following nerve injury proceeds normally in mice lacking DR6

DR6 has been implicated in AxD mechanisms in several neurodegeneration models [6–9]. Given the proposed mechanistic overlap with WD [10], we first sought to confirm that DR6 loss protects axons after injury [5]. The previous work reported that disconnected axons from DR6 null mice are preserved for up to 4 weeks after nerve injury, with a phenotypic penetrance in approximately 38.5% of the animals studied [5]. Quantitative real-time (qRT) PCR and western blot analysis validated successful disruption of DR6 expression in constitutive DR6 null mice (DR6^-/-^), and in another DR6 knockout model with a distinct targeting strategy in which we deleted exon 2 of the *DR6* gene in all tissues including germ cells through Cre-mediated recombination (DR6^-/-, CMV-Cre^) (Fig. S1). Unlike earlier studies using DR6 null mutants [5, 11], we found no significant abnormalities in early developmental myelin formation at postnatal day 1 (Fig. S2), or in g-ratio and axon- and fiber-caliber distributions as assessed by nerve histomorphometry of sciatic and tibial nerve axons from adult DR6^-/-^ and DR6^-/-, CMV-Cre^ animals (Fig. S3). This ensures a normal baseline for our comparative WD studies that follow below.

We transected sciatic nerves of overall 38 DR6^-/-^ and 18 DR6^-/-, CMV-Cre^ mice and assessed axon survival in the distal nerve stump 3 days post-axotomy by light and electron microscopy in comparison to littermate controls. Surprisingly, we found that all studied DR6^-/-^ and DR6^-/-, CMV-Cre^ mice consistently exhibited an almost complete breakdown of axonal structure, identical to control littermate mice (Fig. 1). In contrast, nerve stumps from mice expressing one copy of the *Wld^S^* gene, and mice with loss of the pro-degenerative molecules Phr1/Mycbp2 or SARM1 displayed profound preservation of the vast majority of axons, in line with previous work (Fig. 1).

**Figure 1:**
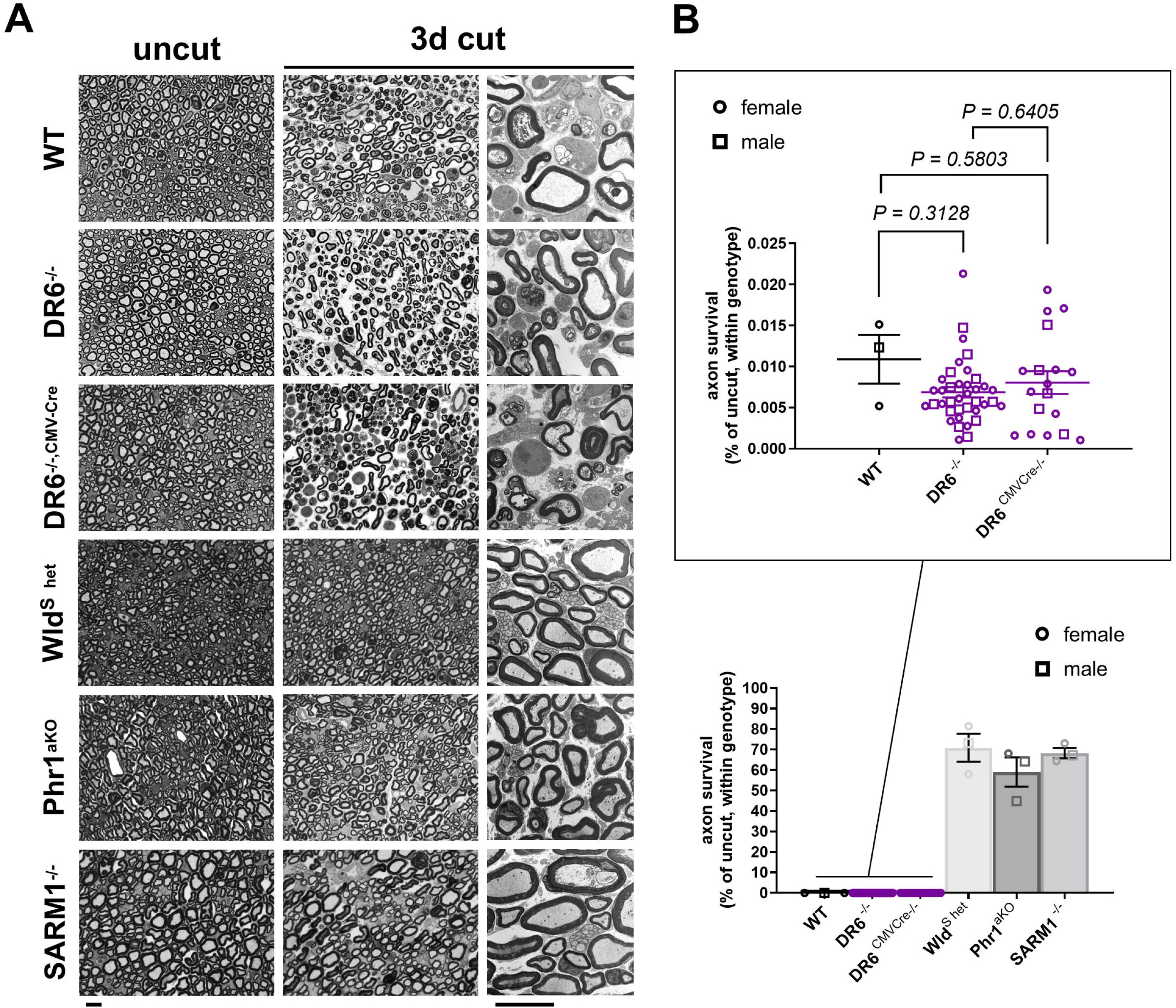
Rapid Wallerian degeneration in mice lacking DR6 in comparison to multiple mutants with delayed degeneration, assessed at 3 days after axotomy. (A) Representative semithin (1^st^ and 2^nd^ columns) and electron micrographs (3^rd^ column) of transverse sciatic nerve sections of distal nerve stumps from wild-type (WT) and the indicated mutant mice 3 days after sciatic nerve transection. Note complete structural disintegration of transected axons with absent or floccular cytoskeleton and collapsed myelin sheaths in the preparations from WT, DR6^-/-^, and DR6^-/-, CMV-Cre^ mice. In contrast, the majority of disconnected axons from heterozygous Wld^S^ mice, and from Phr1 and SARM1 knockout mice, are structurally preserved with uniform cytoskeleton and intact myelin sheaths. Scale bars: 10 µm (B) Quantification of preserved axons in transverse sciatic nerve sections of distal nerve stumps from mice with the indicated genotypes. Each symbol in the scatter dot plots represents the quantification from one animal (% of control axon numbers in micrographs from uninjured contralateral nerve preparations for each animal). All data were obtained from experimental animals between 3 and 12 months of age.

To detect a potentially smaller delay of AxD in the DR6 null mouse models, we next performed lower stringency assays and studied axon survival 30h after nerve injury. This is a time point at which many disconnected axons just start to disintegrate, and ∼50% of myelinated axons in tibial nerves remain structurally intact in wild-type mice as classified on high-resolution light and electron micrographs [4]. Fiber quantification in axotomized nerve segments from DR6^-/-^ and DR6^-/-, CMV-Cre^ mutants demonstrated that the axon survival was statistically indistinguishable from control samples (Fig. 2A). By contrast, Wld^S^ mice, and mutants lacking Phr1/Mycbp2 or SARM1, displayed significantly enhanced axon survival (Fig. 2B).

**Figure 2:**
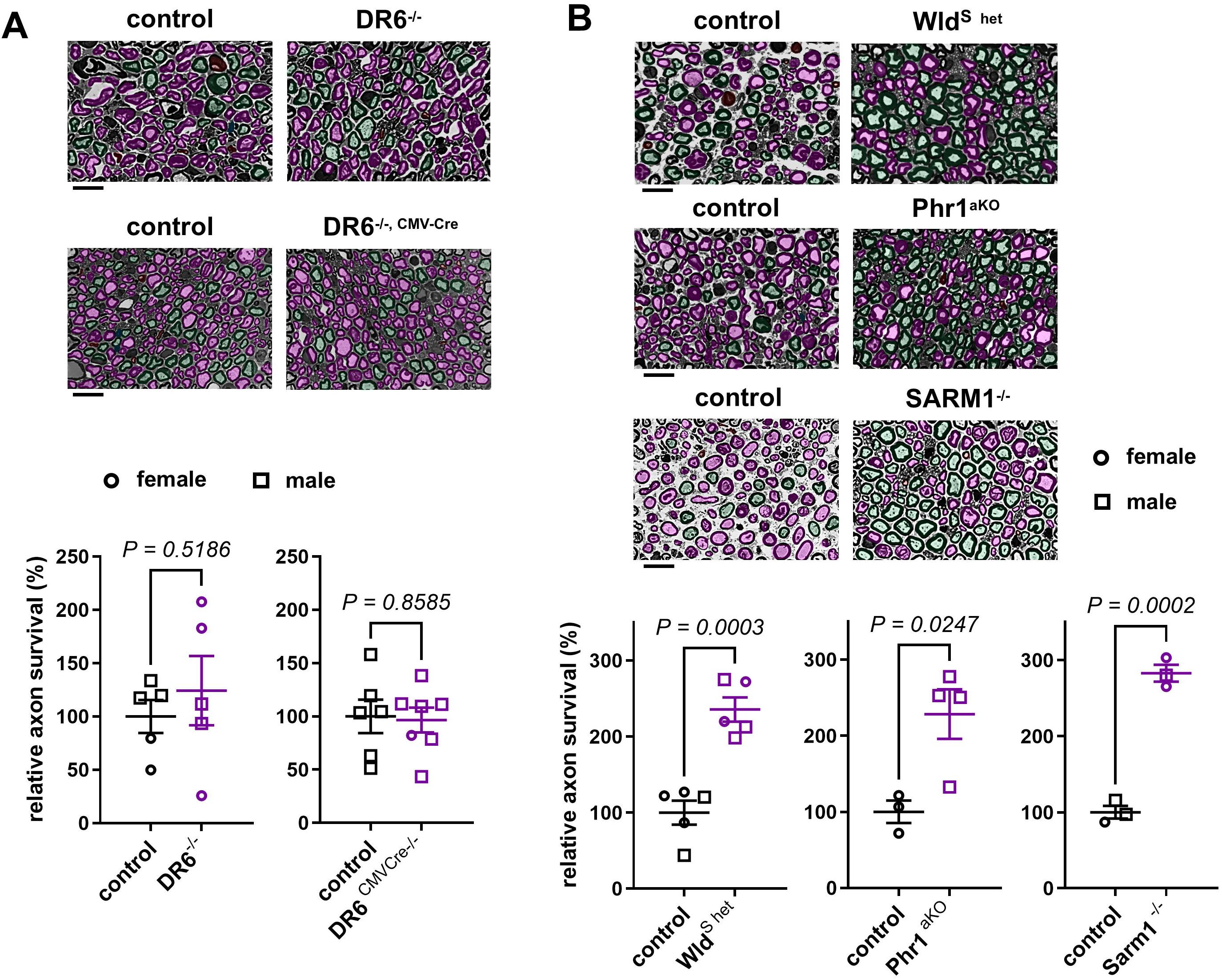
Normal rates of nerve injury-induced axon disintegration in DR6-deficicient mice at earlier stages of Wallerian degeneration. (A) Representative semithin micrographs of transverse tibial nerve sections of distal nerve stumps from 3-months-old mice with the indicated genotypes (top) 30h after sciatic nerve transection with pseudocoloring of intact (turquoise) and structurally degenerated (magenta) myelinated fibers, and corresponding quantifications of relative axon survival (bottom). (B) Representative semithin micrographs of transverse tibial nerve sections of distal nerve stumps from 3-months-old mice with the indicated genotypes (top) 36h after sciatic nerve transection with pseudocoloring of intact (turquoise) and structurally degenerated (magenta) myelinated fibers, and corresponding quantifications of relative axon survival (bottom).Scale bars: 10 µm

We next asked if disconnected nerve stumps from DR6^-/-^ and DR6^-/-, CMV-Cre^ mutants show reduced site-specific phosphorylation (Thr183/Tyr185) and thus activation of c-Jun N-terminal kinases (JNK), as reported previously [5]. Reduced activation of JNKs, members of the mitogen-activated protein kinase (MAPK) family, has been also observed in axotomized optic nerves under conditions of axon protection conferred by SARM1 loss, or expression of a Wld^S^ variant [12]. In accord with the absence of any structural axon protection, we found that the JNK activation in axotomized nerves from DR6^-/-^ and DR6^-/-, CMV-Cre^ mutants was indistinguishable from control preparations (Fig. S4).

Together, these data indicate that the loss of DR6 *in vivo* has no obvious effect on the progression of AxD in axotomized nerves.

### Normal Schwann cell injury responses and myelin remodeling dynamics during WD in the absence of DR6

SCs rapidly transdifferentiate and dismantle their myelin sheaths during WD, a process that is largely orchestrated by the transcription factor c-Jun [13, 14]. The deletion of DR6 has been claimed to alter the typical SC myelin reorganization observed after nerve lesion [5], suggesting abnormal SC transdifferentiation. To determine if DR6-deficient SCs show abnormal reactions to nerve injury, we analyzed c-Jun expression by immunofluorescence on sciatic nerve cross sections distal to the site of nerve injury (Fig. 3). Additionally, we examined the ultrastructure of dedifferentiated SCs (Fig. S5A), and quantified the area occupied by myelin sheaths and myelin debris profiles on osmium tetroxide and toluidine blue stained nerve sections (Fig. S5B, C). We did not detect any significant differences in any of these assays at any time point investigated in DR6^-/-^ and DR6^-/-, CMV-Cre^ samples as compared to control preparations (Fig. 3, S5). These data demonstrate that DR6-deficient SCs show normal injury responses following nerve lesion.

**Figure 3.**
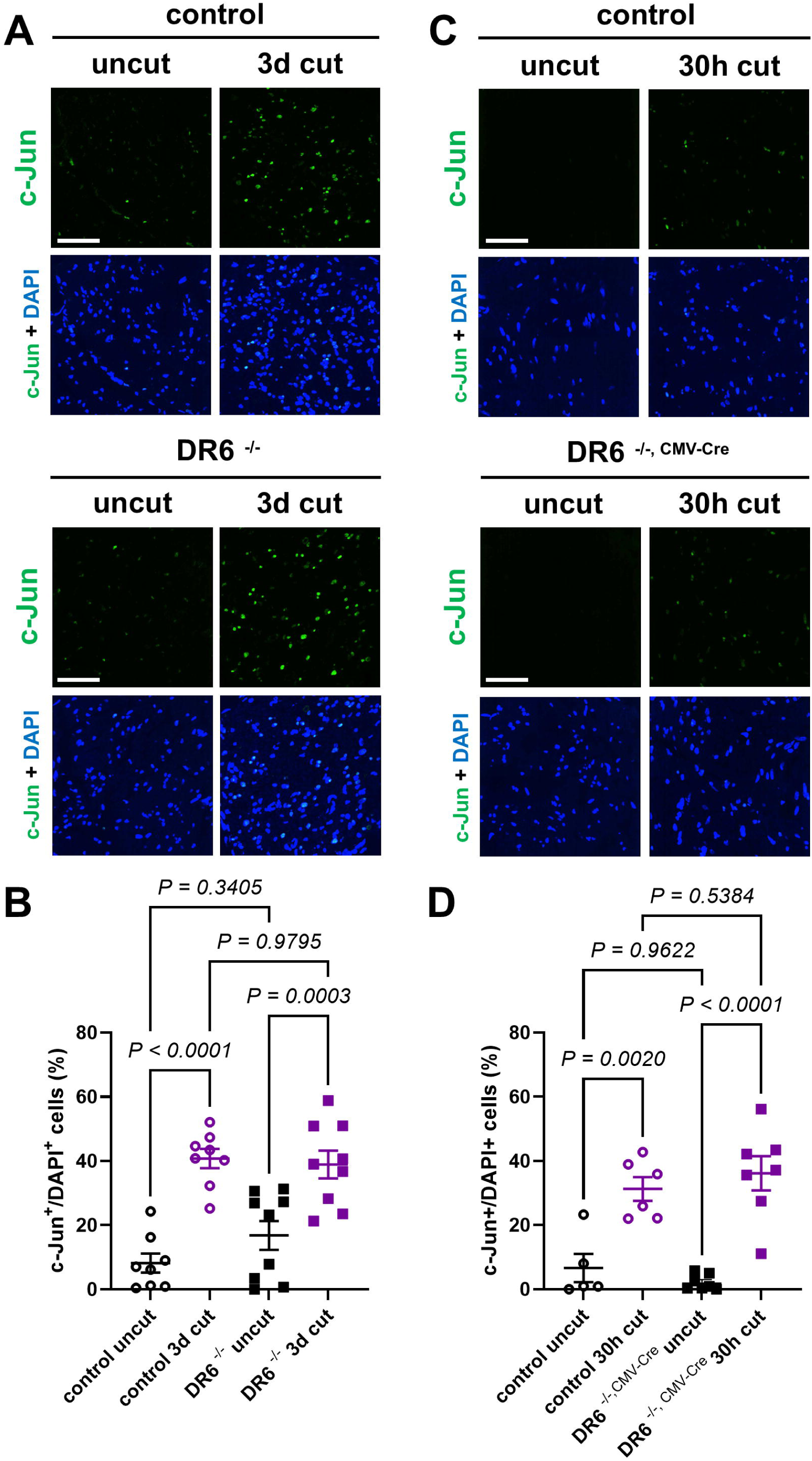
Normal Schwann cell c-Jun injury responses during WD in mice lacking DR6. (A, C) Representative immunofluorescence confocal micrographs for c-Jun with DAPI nuclear counterstaining on transverse frozen sections from contralateral uninjured nerve (uncut) and distal sciatic nerve stumps 3 days and 30 h following axotomy in 3-months-old mice with the indicated genotypes. (B, D) Corresponding quantifications of percentage of c-Jun immunoreactive and DAPI^+^ cells on nerve sections. Scare bars: 50µm.

### Injured DRG neurites in primary neuronal cultures from DR6 knockout mice degenerate rapidly

The results above contrast sharply with the previously reported dramatically delayed AxD during WD in DR6 null mice [5]. In an attempt to reconcile the contradictory findings, we finally considered the possibility that more subtle changes in the stability of injured axons could be potentially unveiled *in vitro* using embryonic neuronal explant cultures from the DR6 mutant models. Such assays were previously extensively used to reveal and characterize axonal DR6 phenotypes, although a key publication in this regard was recently retracted [15]. Importantly, in contrast to the *in vivo* data, no phenotypic penetrance effects in delaying AxD have been reported using *in vitro* assays in the previous DR6 knockout study focusing on WD [5].

We confirmed the loss of DR6 mRNA and protein in dorsal root ganglia explants prepared from DR6^-/-^ embryos, and additionally DR6 *^LoxP/LoxP^* embryos infected with a lentivirus expressing Cre recombinase (Fig. 4A, D, and not shown). However, time-lapse analysis of neurite disintegration after axotomy demonstrated no statistically significant differences between mutant and control preparations from the two models with the only exception of slightly accelerated AxD in the DR6 *^LoxP/LoxP^* preparations with the lentiviral Cre recombinase expression six hours after axotomy (Fig. 4B, C, E, F). This contrasts with the robust axonal protection observed in similar primary culture models devoid of Phr1/Mycbp2, or in cultures with expression of Wld^S^ protein variants we previously reported [16–18]. Together, these findings indicate that the loss of DR6 does not confer resistance to AxD in an established neuronal culture model.

**Figure 4.**
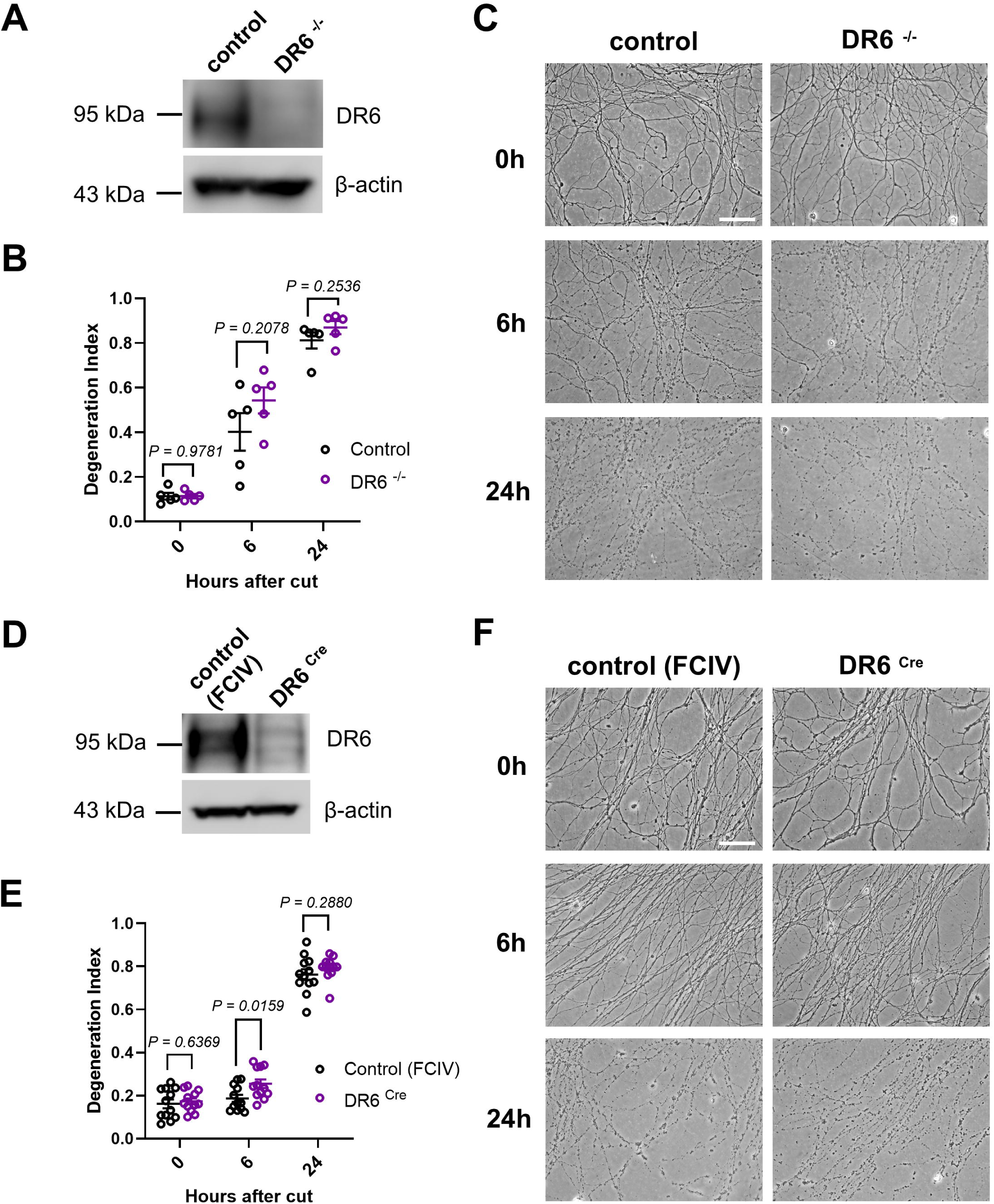
Normal degeneration kinetics of injured DRG neurites in primary neuronal cultures from DR6 knockout mice. (A, D) Western blot analysis (cropped blot images) of DR6 protein expression in dorsal root ganglia (DRGs) from DR6^-/-^ embryos and from DR6 *^LoxP/LoxP^* embryos infected with a lentivirus expressing Cre recombinase (DR6 ^Cre^), together with the respective control preparations. (B, E) Time course of neurite fragmentation, quantified as degeneration index (DI) (see Experimental Procedures) in DRG preparations from embryos with the indicated genotypes or lentiviral infection conditions. The data points shown in (B) represent the averaged neurite DI values calculated from multiple DRG preparations from each embryo at the indicated time point after injury (one data point = one embryo). The data points shown in (E) represent the averaged neurite DI values calculated from multiple micrographs acquired from each DRG preparation in a cell culture well at the indicated time point after injury (one data point = one well). (C, F) Representative phase-contrast micrographs of disconnected DRG neurites from embryos with the indicated genotypes and lentiviral infection conditions at the indicated time points after axotomy. Scale bars: 50µm.

## Discussion

The main motivation for this study was to determine if the glial injury responses previously suggested as abnormal in DR6-deficient mice play a role in the extended axon survival observed during WD [5]. Unexpectedly, despite using the same DR6 mutant mouse line as the original study, we did not observe delayed AxD or the aberrant myelin remodeling described in that work. Our findings demonstrate that DR6 does not function as a pro-degenerative component within the axonal auto-destruction pathway, nor does it play a significant role in regulating the glial injury responses and myelin remodeling dynamics characteristic of WD in the peripheral nervous system (PNS). Our study underscores the importance of rigorous validation when assigning functional relevance to molecules with possible roles in the WD pathway.

The axon with its associated glial cells forms a specialized structural and functional unit that enables long-range neural communication. Much like how compromised cells initiate self-destruction, injured axons undergo a related but molecularly distinct process during WD, characterized by rapid metabolic collapse [12, 19]. We recently showed that SCs modulate this process through early injury responses that enhance metabolic coupling with axons [4].

Importantly, the WD pathway of axonal self-destruction is believed to contribute significantly to the progression of various neurodegenerative diseases, highlighting both its cell-autonomous and non-cell-autonomous suppression as promising therapeutic strategies. To date, apart from expression of the aberrant fusion protein Wld^S^, only four endogenous molecules – Nmnat2, SARM1, Phr1/Mycbp2, and DR6 - have been implicated to play a significant role in the WD pathway in mammals [5, 16, 20, 21]. Their overexpression or inactivation has been reported to confer drastic resistance to experimentally-induced WD, with disconnected axons in the PNS surviving for days to weeks, unlike wild-type axons, which fully degenerate within 48–72 hours on structural level. While Wld^S^, Nmnat2, SARM1, and Phr1/Mycbp2 are cell-autonomously involved in the common WD pathway leading to axonal metabolic failure, DR6 has emerged as a more elusive candidate. Initial reports suggested that DR6, along with its putative ligand amyloid precursor protein (APP), mediates developmental axon pruning [22–24]. However, this interpretation has been complicated by a recent retraction [15] and conflicting findings [25]. A later study proposed a striking new role for DR6 in WD, reporting that severed axons in DR6 knockout mice remained structurally preserved for up to four weeks post-injury even though separated from their parent neuronal cell bodies [5]. However, this phenotype was observed in fewer than half the animals, suggesting incomplete penetrance. The axonal phenotype, when present, was similar to that in Wld^S^ mice, which also showed incomplete phenotypic penetrance in this study [5]. Moreover, it was found that the subset of DR6-deficient mice with axon protection displayed abnormal patterns of myelin remodeling during delayed WD, suggesting aberrant SC injury responses that could have contributed to the axonal phenotype. Furthermore, another study reported that neuronal DR6 expression regulates SC proliferation and inhibits developmental myelin formation in SCs [11]. These findings together raised the possibility that inactivation of DR6 might influence both neuronal and glial functions and their crosstalk during WD. However, to date, no subsequent studies have expanded upon or validated these discoveries.

In surprising contrast to the earlier data, our results provide compelling evidence that DR6 does not regulate AxD or glial injury responses during WD in the PNS. We employed a robust experimental design and have extensively characterized the kinetics of WD in two independent DR6 mouse knockout models using a high number of experimental animals and multiple *in vivo* and *in vitro* assays. For the first time, we present a comprehensive side-by-side comparison of key mouse mutants that have previously been shown to exhibit markedly delayed WD following nerve injury. We found no evidence that loss of DR6 expression, as confirmed by qRT-PCR and western blotting, antagonizes AxD during WD. Even under low-stringency conditions designed to detect subtle axonal protection, axon death progressed normally. These findings align with a prior electrophysiological study in the central nervous system, which suggested normal degeneration of optic nerve axons in DR6-deficient mice [26]. We also did not find any changes in the SC injury responses and myelin reorganization known to be orchestrated by the transcription factor c-Jun during WD in the PNS [13, 14] in our DR6 mutant models at two post-injury time points that are frequently used to assess c-Jun expression and SC dedifferentiation characteristics We therefore did not examine SC injury responses and myelin reorganization at 14 days after nerve injury, as performed in the previous study [5], because no evidence of axonal protection was seen at earlier post-lesion stages.

The prominent discrepancy between our results and the prior study [5] remains unresolved. One possibility is that a spontaneous germline mutation arose within the previously used mouse colony, conferring axonal protection in a subset of animals through a mechanism unrelated to DR6. Such mutation, perhaps conceptually similar to the one responsible for the Wld^S^ phenotype that occurred spontaneously in the Olac breeding colony in England [27], could then have propagated within the colony via autosomal dominant inheritance. Although we obtained our DR6^-/-^ breeders directly from the laboratory that initially reported the protective phenotype [5], it is conceivable that the specific mice we received did not carry the hypothetical second-site mutation. Alternatively, the phenotypic discrepancies may reflect the influences of unknown genetic modifiers, epigenetic factors, environmental conditions, or age-related differences that could affect the WD pathway. These variables could potentially explain the reported partial penetrance and the previously described developmental anomalies in DR6 null nerves [5, 11].

We also considered if changes in the nerve immune microenvironment might account for the discrepancies. For example, altered macrophage recruitment or activation, which is known to influence axon integrity, could contribute to differences in WD dynamics [28]. However, despite these possibilities, we note that we have never observed incomplete phenotypic penetrance effects in Wld^S^ mice we developed [17, 18, 29–31] or in mutants lacking SARM1 or Phr1/Mycbp2 [16], either in our hands or in reports from other laboratories over the last decades. This supports the view that the manipulation of these core components of an evolutionary conserved pathway produces fully penetrant protective phenotypes. Nonetheless, environmental modulation of axon protection in Wld^S^ mice has been described, although the reported effects are relatively modest [32].

DR6 inhibition has shown therapeutic benefit in several models of neurodegeneration, including amyotrophic lateral sclerosis [7, 8], multiple sclerosis [33, 34], prion disease [9], and *in vitro* amyloid-beta toxicity [6]. However, no protective effects were observed in two separate mouse models of Alzheimer’s disease [24]. While these effects have often been attributed to reduced neuronal death [6, 8], changes in immune function [34], or enhanced myelination [33], the suggestion that DR6 suppression stabilizes injured axons raised the possibility that axon protection itself may have contributed to disease amelioration. Our data argue against this interpretation in the context of WD. Instead, as mentioned earlier, DR6 may play a regulatory role in alternative forms of AxD, such as axon pruning, which is known to shape and refine neural circuits during development and may also be reactivated under pathological conditions [35, 36]. Future studies using disease-specific models will be necessary to explore this hypothesis.

## Author contributions

E.B. and B.B. conceived and supervised the study. E.B. and B.B. designed and planned the experiments. E.B., H.H., and B.B. performed the experiments and analyzed and interpreted the data shown in the figures. B.B. and E.B. wrote the manuscript.

## Acknowledgements

We thank Genentech and Christopher Deppmann (University of Virginia) for providing the constitutive and conditional DR6 mutant mice, Konstantinos Tsesmelis for technical experimental assistance, Ekaterina Stepanova and Alessio Colombo for genotyping information, Devin Donich for help with the g-ratio measurements, and Sara Bombardelli and the University Laboratory and Animal Resources (ULAR) at The Ohio State University College of Medicine for mouse husbandry assistance. This study was supported by the NIH-NINDS grant R01NS123450 (to E.B. and B.B.).

## Materials and Methods

### Mice

All experiments were performed in compliance with the Association for Assessment of Laboratory Animal Care policies and approved by The Ohio State University College of Medicine Animal Care and Use Committee. Mice were housed under specific pathogen-free conditions at 70 °F, 50% room humidity, 12-h light/12-h dark cycle and received ad libitum access to water and food. Mice from different genotypes were group-housed in separate cages. The mice used in this study did not undergo any procedures before their stated use. The mice were of mixed sexes. In previous studies we found no significant differences in the rates of injury-induced axon degeneration between female and male mice. Mice within individual comparative *in vivo* experiments were littermates or, if use of littermates was not possible because of litter sizes not large enough, mice from different litters were age--matched for comparative experiments. Controls were regarded as littermates or age--matched mice from different litters carrying wild-type alleles or floxed alleles with no Cre recombinase expression for the respective genotype groups. Constitutive DR6 knockout mice (DR6^-/-^) with deletion of exons 2 and 3 (on C57BL/6J.129S mixed background) and conditional DR6 mice with exon 2 flanked by loxP sites (DR6 *^LoxP/LoxP^*) were generated by Genentech [37, 38] (genetic background unknown). To generate DR6^-/-, CMV-Cre^ mice, we crossed DR6 *^LoxP/LoxP^* mice to CMV-Cre transgenic mice (strain no. 006054, The Jackson Laboratory, made congenic on C57Bl/6J background). For this study, we additionally used heterozygous Wld^S^ [29], Phr1 aKO [16], and SARM1^-/-^ mice (strain no. 018069, The Jackson Laboratory, made congenic on C57Bl/6J background). Genotyping was performed by PCR strategies using standard procedures and appropriate primers (sequences available upon request).

### Multiplex TaqMan quantitative real-time PCR

Brain tissue from DR6^-/-^ and DR6^-/-, CMV-Cre^ mutants and the respective control mice were homogenized using a Bullet Blender (Next Advance, model BBX24B) and RNAase-free zirconium oxide beads (0.5mm diameter, Next Advance). Total brain RNA was extracted using PureLink RNA Mini Kit (Thermo Fisher Scientific) according to the manufacturer’s guidelines, and RNA concentration and purity were assessed using a NanoDrop One spectrophotometer (Thermo Fisher Scientific). To eliminate potential genomic DNA contamination, 4 µg of total RNA per sample were treated with EzDNAase (Invitrogen) according to the manufacturer’s instructions. The resulting RNA was split into two aliquots: 2 µg were reverse transcribed into cDNA using SuperScript VILO Master Mix (Thermo Fisher Scientific), and the remaining 2 µg underwent the same reaction without the reverse transcriptase enzyme to generate a no-RT control. Reaction times and volumes followed the manufacturer’s guidelines. DR6 expression was tested with TaqMan gene expression predesigned assay Mm00446361_m1 with PAM-MGB probe [6], and the housekeeping gene peptidyl-prolyl isomerase A (PPIA) was tested with TaqMan gene expression assay ID Mm02342430_g1 (Thermo Fisher Scientific) with VIC-MGP probe. Briefly, in each well of a 96-well plate (MicroAmp Optical 96-well reaction plate with barcode, Applied Biosystems), 1µl of cDNA was tested with 9 µl of Mastermix containing 0.5µL of TaqMan probe, using a QuantStudio 3 Real Time PCR instrument (Applied Biosystems). The thermal-cycling conditions included an Uracil-N glycosylase incubation step (50°C for 2 minutes), followed by Polymerase activation (95°C for 20 seconds), and 40 cycles of PCR reaction (denaturation at 95°C for 3 seconds and annealing/extension at 60°C for 30 seconds). ROX reference reading (passive reference) and Cycle-Threshold (CT) values were collected and visualized with QuantStudio 3 software and analyzed with the DeltaDeltaCT method. DR6 expression levels were calculated and normalized to PPIA levels in the same well of the plate.

### Western blot analysis

Mouse brains were quickly dissected and homogenized using a Polytron homogenizer (PT 1200C), followed by sonication with a Qsonica sonicator (Q500) equipped with a microtip. Sciatic nerves were quickly dissected, the epineurium removed in ice-cold phosphate buffered saline (PBS), and directly sonicated as above. Both brains and nerves were homogenized in RIPA buffer supplemented with protease and phosphatase inhibitors. Embryonic dorsal root ganglia preparations were washed with PBS to remove culture medium and then manually collected with forceps, placed in a tube with RIPA buffer supplemented with protease and phosphatase inhibitors, and then vortexed. After lysis, samples were spun at 10,000g for 10 minutes at 4°C to remove insolubilized material. Western blotting of tissue lysates was performed using Bolt Mini gels and Mini Blot wet transfer modules (Invitrogen) according to the manufacturer’s instructions. For detection of the DR6 protein, tissue and cell homogenates were processed under denaturing conditions on Tris-Acetate gels according to the manufacturer’s protocol optimized for large molecular weight proteins (Thermo Fisher Scientific). For detection of JNK and p-JNK (Thr183/Tyr185) in nerve lysates, proteins were separated on 4-12% Bis-Tris gradient gels (Thermo Fisher Scientific). The following antibodies were used in a solution of 5% bovine serum albumin (BSA) in Tris-buffered saline (TBS) with 0.1% of Tween20: anti-DR6 clone 6B6 (1:500, catalog no. MABC1594, Millipore Sigma), anti-phospho-SAPK/JNK (Thr183/Tyr185) clone 81E11 (1:1000, catalog no. 4668, Cell Signaling), anti-SAPK/JNK (1:1000, catalog no. 9252, Cell Signaling), anti-b-actin (1:10,000, catalog no. A5316, Millipore Sigma). Horseradish peroxidase-conjugated secondary antibodies from Cell Signaling Technology and Jackson ImmunoResearch were used for signal detection. Blot documentation was performed with a G:Box mini 6 digital imaging system (Syngene). The integrated band intensities of protein bands were determined using ImageJ software to calculate p-JNK(Thr183/Tyr185)/JNK ratios.

### Histomorphometry of intact nerves

The embedding of nerve samples in Araldite 502 epoxy resin (Polysciences) and subsequent cutting and staining of 500-nm semithin cross sections was performed as described previously [4, 39, 40]. High-resolution tile scans of the entire sciatic or tibial nerve cross sections were acquired with a DMi8 digital imaging system equipped with a 100x high-numerical aperture objective and oil immersion condenser using the Application Suite X 3.7.2 software (Leica Microsystems). The quantification of thinly myelinated axons on entire sciatic nerves from P1 mouse pups was performed manually using the ImageJ Cell Counter plug-in. The *g* ratios of individual myelinated axons in sciatic or tibial nerves as a measure of myelin thickness were determined using a plug-in for the ImageJ software that allows semiautomated analysis of randomly selected axons on nerve transverse sections (http://gratio.efil.de). One hundred randomly chosen fibers were measured per mouse nerve. Cumulative *g* ratios were calculated for each mouse by averaging all individual *g* ratios. Axon and fiber caliber distributions were quantified from sciatic or tibial nerve micrographs using ImageJ. The Wand tracing tool was used to manually delineate individual regions of interest (ROIs). For axon caliber measurements, the tool was applied to select the axonal area (excluding the myelin sheath), with the tolerance adjusted to accurately define each axon’s boundary. For fiber caliber measurements, the entire fiber (axon plus myelin sheath) was selected using the same approach. The resulting ROI data were exported to Microsoft Excel, and calibers were calculated using the formula A = π (d/2)^2, where A is the area and d is the diameter. Binned distributions of axon and fiber calibers were calculated and visualized using GraphPad Prism software. All quantifications were conducted blinded to the genotypes of the animals.

### Electron microscopy

Transmission electron microscopy was carried out as previously described [4, 39]. Electron micrographs of randomly selected areas on 85 µm thick ultrathin nerve cross sections stained with uranyl acetate and lead citrate were taken with a FEI Tecnai transmission electron microscope.

### Unilateral sciatic nerve transection

Mice were deeply anesthetized using the SomnoSuite Low-Flow Anesthesia delivery system (Kent Scientific) operated with isoflurane. Right sciatic nerves were exposed and transected with surgical micro-scissors close to the sciatic notch with the contralateral nerve serving as the control. The wound was closed with surgical thread or clips and buprenorphine was administered as a postsurgery analgesic. Upon nerve removal from humanely killed mice, the lesion site was inspected to verify complete transection. Distal sciatic and tibial nerve stumps and contralateral uninjured nerve segments were processed for semithin/electron microscopy, or for immunofluorescence as described below.

### Analysis of axonal survival in axotomized nerve stumps

Axon survival after unilateral sciatic nerve transection was determined using highly established methods for the evaluation of axonal integrity [4, 18, 29, 30]. Axon survival analysis 3 days post-axotomy was performed on semithin micrographs from randomly selected areas on sciatic nerve cross sections acquired with a DMi8 digital imaging system equipped with a 100x high-numerical aperture objective. One micrograph per mouse nerve was quantified. On each micrograph, the number of all structurally intact myelinated axons was manually counted using the Cell counter plug-in of the ImageJ software. Axon survival was expressed as percentage of intact axons relative to the total number of intact myelinated axons in corresponding uninjured control nerve micrographs. Axon survival at 30-36h after axotomy was assessed on montaged tile scan images of entire tibial nerves, acquired with a Leica DMi8 digital imaging system as described above, and similar to our previously published work [4]. Because Wallerian degeneration at such early stages following axotomy is asynchronous resulting in substantial heterogeneity of axons disintegration within a nerve [41], this method prevents the sampling bias that is associated with quantification of axons from individual micrographs taken from select areas of the nerve. Criteria for structurally intact axons after axotomy were uniform axoplasm with presence of non-swollen mitochondria and normal myelin sheaths. Criteria for degenerated axons were degraded axoplasm, absence of mitochondria and collapsed myelin sheaths. All structurally intact and all degenerated myelinated axons were counted on each tibial nerve montage, and their ratio was determined for each experimental animal (relative axon survival). Scoring was documented with the ImageJ software using the Cell counter plug-in. The experimenter was blind to the genotypes of the mice during data acquisition. Relative axonal survival is reported as the percentage of the respective control group.

### Analysis of myelin remodeling in axotomized nerve stumps

Quantification of the area occupied by myelin (sheaths and myelin debris) in transverse semithin sections of axotomized sciatic and tibial nerves was performed using ImageJ. This analysis leverages the high contrast provided by osmium tetroxide and toluidine blue staining of lipid-rich myelin, which enables reliable thresholding and binarization of myelin structures on nerve cross sections. The binarized images were used to calculate the proportion of the nerve cross-sectional area occupied by myelin. We used axotomized nerves from mice lacking c-Jun in Schwann cells (c-Jun *^loxP/loxP^*, P0^Cre^) with suppressed Schwann cell transdifferentiation as a positive reference which show abnormal myelin remodeling and clearance during Wallerian degeneration. This results in significantly larger myelin areas occupied during Wallerian degeneration as compared to control preparations.

### c-Jun immunofluorescence and quantification

Segments of control and injured sciatic nerves were immersion fixed in 4% paraformaldehyde/0.1M PBS for 1h at 4°C, washed in PBS, cryoprotected in 30% sucrose, embedded in O.C.T. compound, and sectioned at 12 µm on a Leica cryostat (Leica Biosystems). The cross sections on adhesive glass slides were washed with TBS, permeabilized with 0.25% Triton X-100 and 0.5M NH_4_Cl in 0.05M TBS for 10 min, blocked with 5% goat serum in 0.05M TBS for 1 h, and then incubated with anti-c-Jun antibody (1:200, catalog no. 9165, Cell Signaling Technology) diluted in 5% goat serum/0.05M TBS overnight at 4 °C. After washing in TBS, anti-rabbit secondary antibody coupled to Alexa Fluor 488 (1:500, catalog no. 115-545-003) was applied for 1 h, the sections counterstained with DAPI (1:10,000, catalog no. 62247, Thermo Fisher Scientific) and mounted in VECTASHIELD Antifade Mounting Medium. Micrographs of the stained cross sections were captured with an Andor Dragonfly 200 spinning disc confocal imaging system (Oxford Instruments) and a 63x high numerical aperture objective. Adjustments of brightness and contrast were applied with ImageJ and Microsoft Powerpoint equally across the entire image and were applied equally to control preparations for all the data presented. Two representative maximum intensity z-projection images from randomly selected equal volumes were taken per nerve cross section and used for the subsequent quantification. The quantification of c-Jun^+^/ DAPI^+^ cells versus c-Jun^-^/DAPI^+^ cells used to calculate the percentage of cells showing c-Jun immunoreactivity was carried out blind to the mouse genotypes.

### Preparation of mouse embryonic dorsal root ganglia cultures

Embryonic dorsal root ganglia (eDRGs) were isolated from mouse E13.5 embryos. DR6^+/-^females were timed-mated with DR6^+/-^ males, and embryos from each litter were processed individually and genotyped after eDRG plating. eDRGs from DR6^+/+^ embryos were used as control preparations. DR6 *^loxP/loxP^* females were time-mated with DR6 *^loxP/loxP^* males, and all extracted embryos from these litters were pooled together for eDRG isolation. eDRGs were dissociated and plated as previously described [42]. A suspension of about 25,000 cells/µl was plated as a 2µl droplet into a well within a 24-well plate. Neurons were cultured on poly-D-lysin and laminin coated wells in Neurobasal medium supplemented with Glutamine (2mM), Penicillin/Streptomycin, serum-free B27, nerve growth factor (NGF, 50ng/ml), and the anti-mitotic agents uridine and 5-Fluoro-2’-deoxyuridine (1µM) to inhibit non-neuronal cell proliferation. At 2 days *in-vitro* (DIV 2), neurons from DR6 *^loxP/loxP^* embryos were infected with either a control lentivirus (FCIV), or a lentivirus expressing Cre recombinase (CRE), both based on the same vector backbone and co-expressing EGFP, as previously described [16]. Infection efficiency was monitored via fluorescence microscopy based on EGFP expression.

### Analysis of *in vitro* axon degeneration

At DIV 7, when eDRG neurites had developed an extended radial arborization pattern, neurites were transected from their neuronal cell bodies using a microblade under microscopic guidance. The region distal to the transection site was imaged by phase contrast at the indicated time points with a Leica DMi8 digital imaging system and processed as previously described [42]. The micrographs were analyzed in ImageJ using a previously published degeneration index (DI) macro [43]. This macro binarizes the images and calculates the area occupied by fragmented neurites, normalized to the total neurite area. The resulting DI value models the extent of structural axon disintegration during Wallerian degeneration, with values above 0.8 reflecting complete axonal disintegration. For the comparison of DR6^+/+^ and DR6^-/-^ embryos, the time-course of neurite fragmentation for each embryo was obtained by averaging the DI values calculated from multiple phase contrast micrographs taken from each cell culture well at each time point. For the comparison of lentiviral infected neurons from DR6 *^loxP/loxP^* embryos, the time-course of neurite fragmentation in each eDRG cell culture well preparation was obtained by averaging the DI values calculated from multiple phase contrast micrographs taken from each cell culture well.

### Statistics and reproducibility

All shown micrographs (semithin and electron microscopy, immunofluorescence, phase contrast microscopy) are representative of at least three biological replicates (mice or cell culture preparations). All statistical analyses and data visualization were performed using Graphpad Prism software. An unpaired, two-tailed Student’s t-test was used to compare two groups. A two-way analysis of variance (ANOVA) with Sidak’s multiple comparisons test was used to compare more than two groups. Data are presented as arithmetic mean ± standard error of the mean (SEM). Statistical differences were considered to be significant when *P* < 0.05.

---

**Figure S1: Disrupted DR6 expression in two DR6 knockout mouse models**

(A, C) Quantitative real-time PCR analysis of relative brain DR6 mRNA levels from control and mutant mice with the indicated genotypes. Each dot represents the measurement from one mouse.

(B, D) Western blot analysis (cropped blot images) of brain lysates from control and mutant mice with the indicated genotypes probed with the shown antibodies. Each western blot lane represents the brain lysate data from one individual mouse.

**Figure S2: Normal developmental myelination in sciatic nerves from mice lacking DR6 at postnatal day 1**

(A, C) Representative semithin micrographs (A, C) of transverse sciatic nerve sections from control and mutant mouse pups at postnatal day 1 with the indicated genotypes. Red arrows depict examples of axons with nascent myelination. Scale bars: 10µm.

(B, D) Quantification of myelinated axons in transverse sciatic nerve sections from control and mutant pups at postnatal day 1 with the indicated genotypes. Each dot represents the quantification obtained from one mouse pup.

**Figure S3: Normal nerve histomorphometry in DR6-deficient mice**

(A, C) Quantification of *g* ratios (left: scatter plots show *g* ratios of individual myelinated axons as function of axon diameter, right: corresponding cumulative *g* ratios per animal) in tibial (A) and sciatic nerves (C) from 3-months-old control and mutant mice with the indicated genotypes. (B, D) Axon caliber and fiber caliber (axon plus myelin sheath) distributions in tibial (B) and sciatic nerves (D) from 3-months-old control and mutant mice with the indicated genotypes.

Each dot represents the measurement from one mouse.

**Figure S4: Normal JNK activation in sciatic nerves from DR6-deficient mice**

(A, C) Representative western blots (cropped blot images) of lysates from uninjured (0 min) and axotomized distal sciatic nerve stumps (30 min after axotomy) from 1-2-months-old control and mutant mice with the indicated genotypes probed with the shown antibodies. Each lane represents the result from one mouse sciatic nerve.

(B, D) Densitometric quantifications with statistical analysis of western blot data for all experimental animals used. Each dot represents a measurement from sciatic nerve lysate from one mouse.

**Figure S5: Normal ultrastructure of transdifferentiated Schwann cells and normal myelin remodeling dynamics during Wallerian degeneration in DR6-deficient mice**

(A) Representative electron micrographs of transdifferentiated Schwann cells (red arrowheads) that lost axonal contact from distal sciatic nerve stumps 3 days following nerve transection in 3-months-old control and DR6 knockout mice with the shown genotypes. Note similar ultrastructure with marked cytoplasmic expansion, increased organelle content, and myelin debris (myelin ovoids) in Schwann cell bodies in all genotypes. Scale bar: 2µm.

(B, C) Quantification of area occupied by myelin sheaths and myelin debris in transverse nerve sections from 3-months-old control and mutant mice with the indicated genotypes. Each dot represents the quantification obtained from one nerve.

