## Supplementary figures and images for "Death receptor 6 does not regulate axon degeneration and Schwann cell injury responses during Wallerian degeneration"

### Figure S1

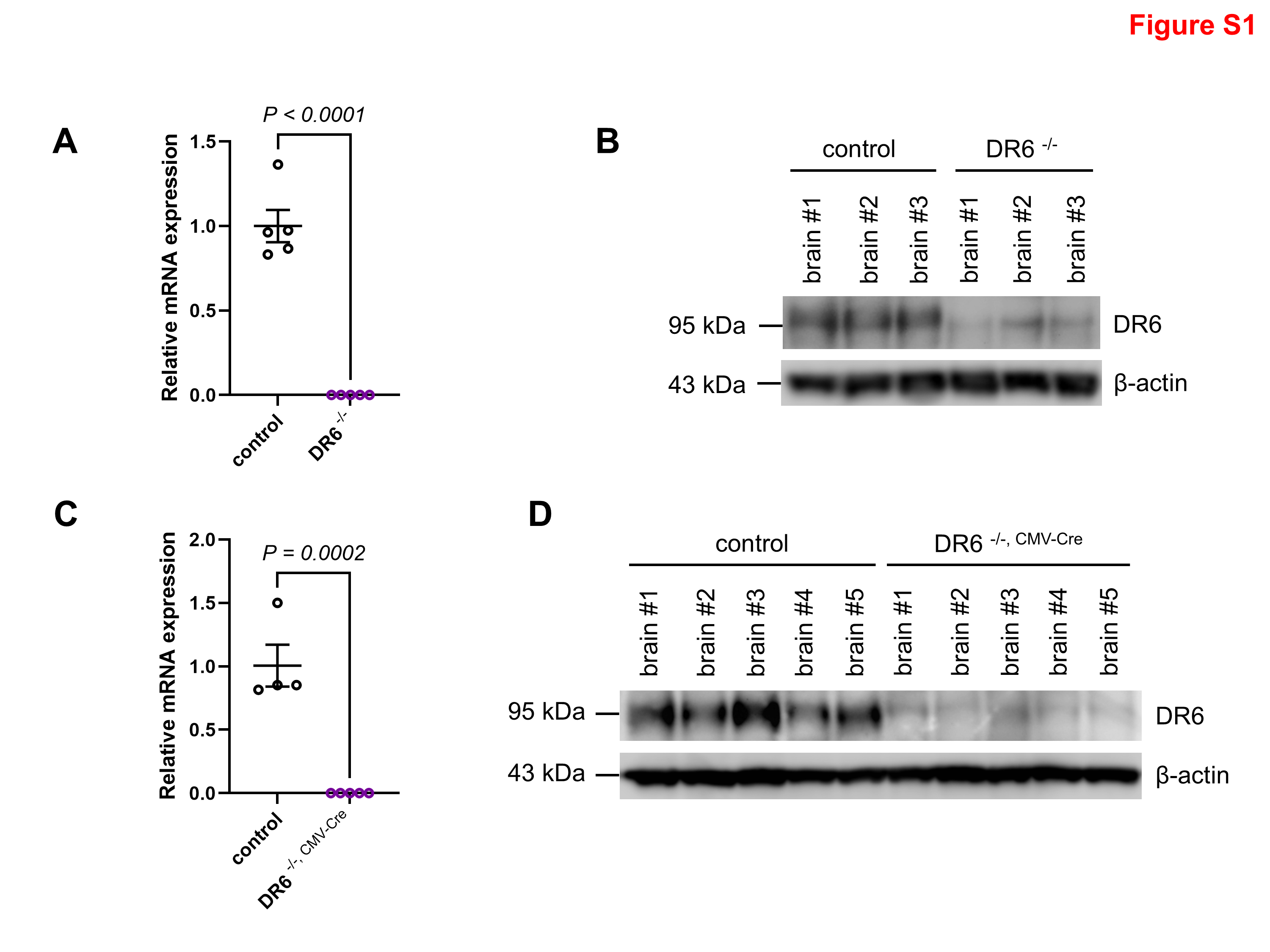

### Figure S2

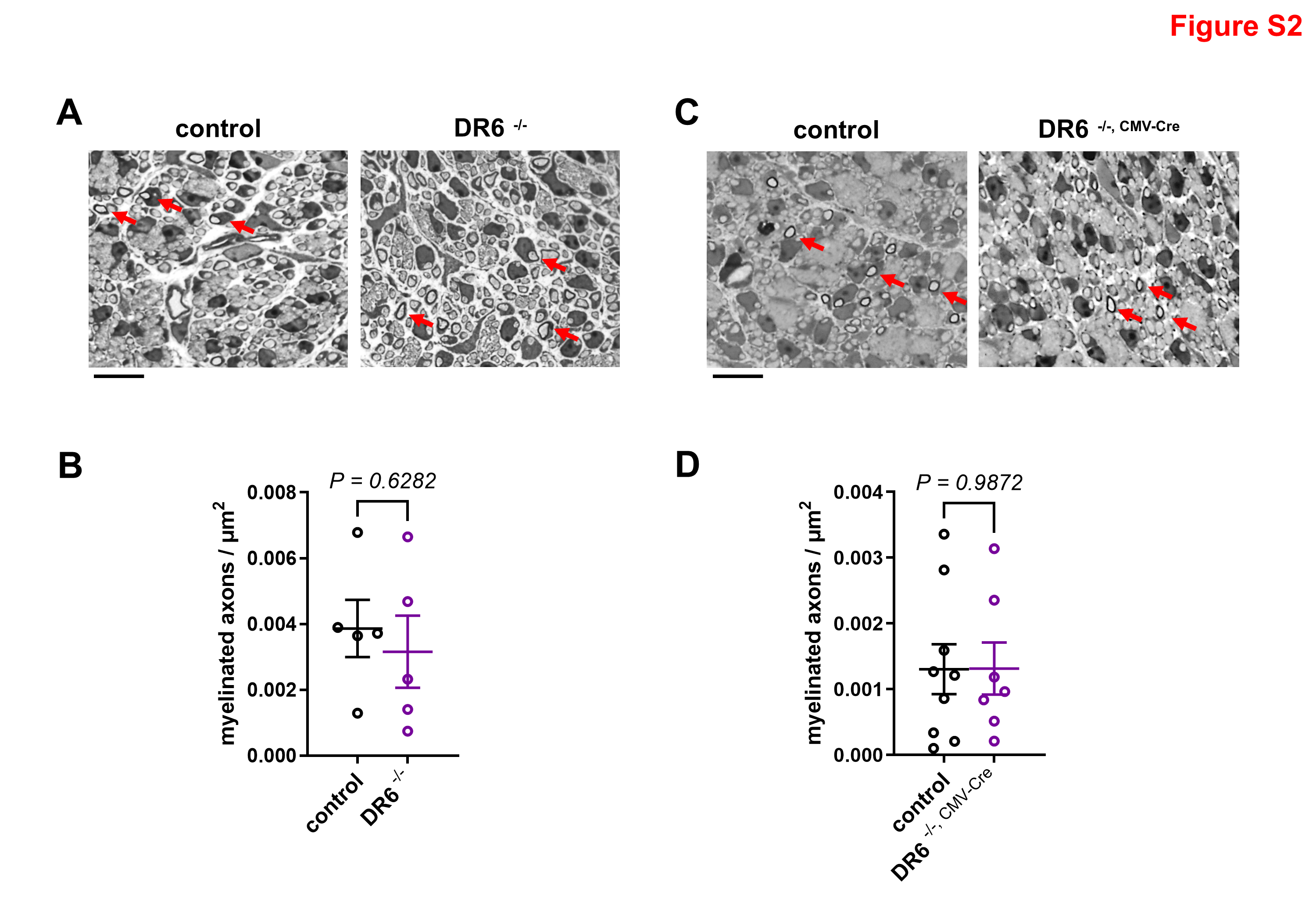

### Figure S3

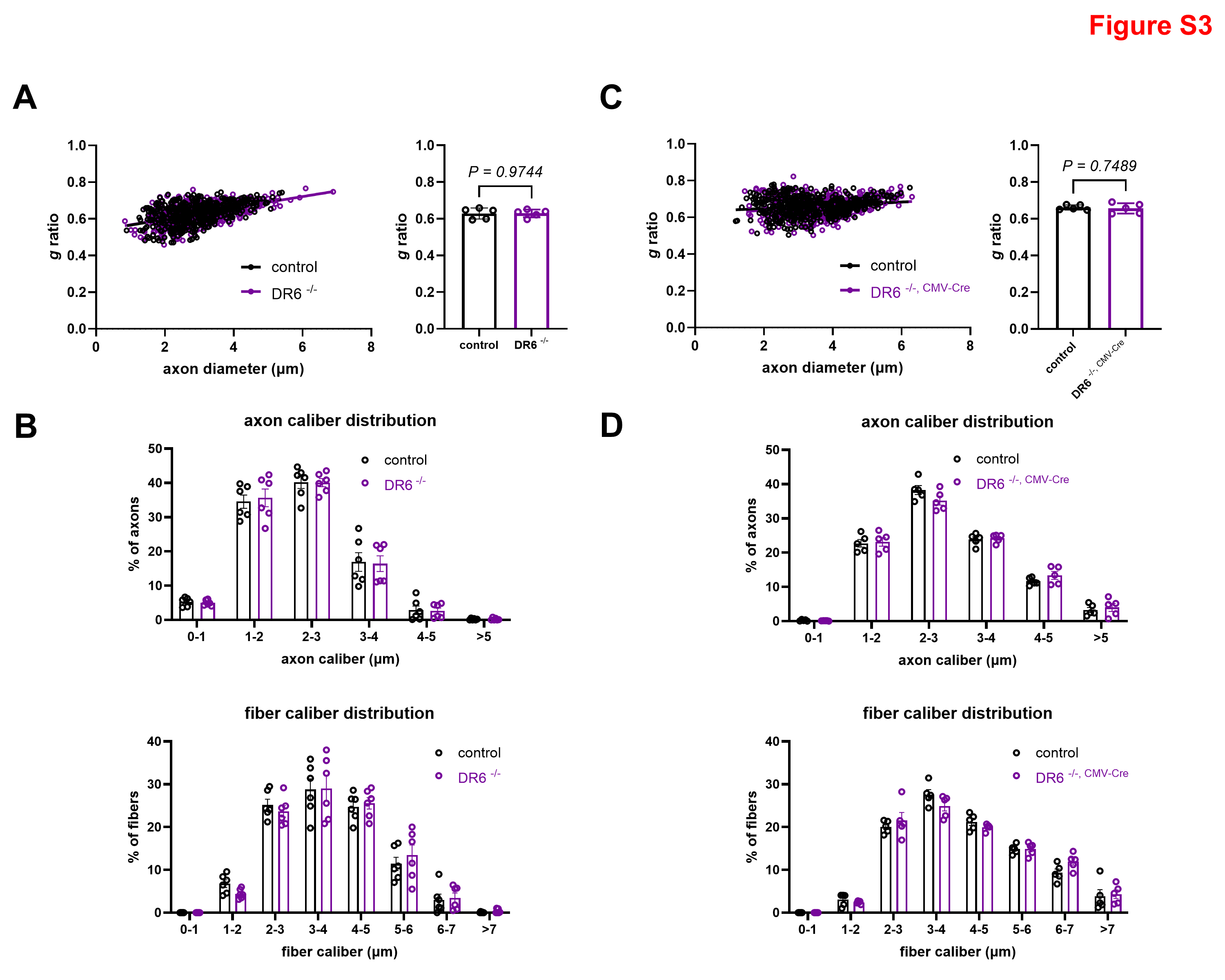

### Figure S4

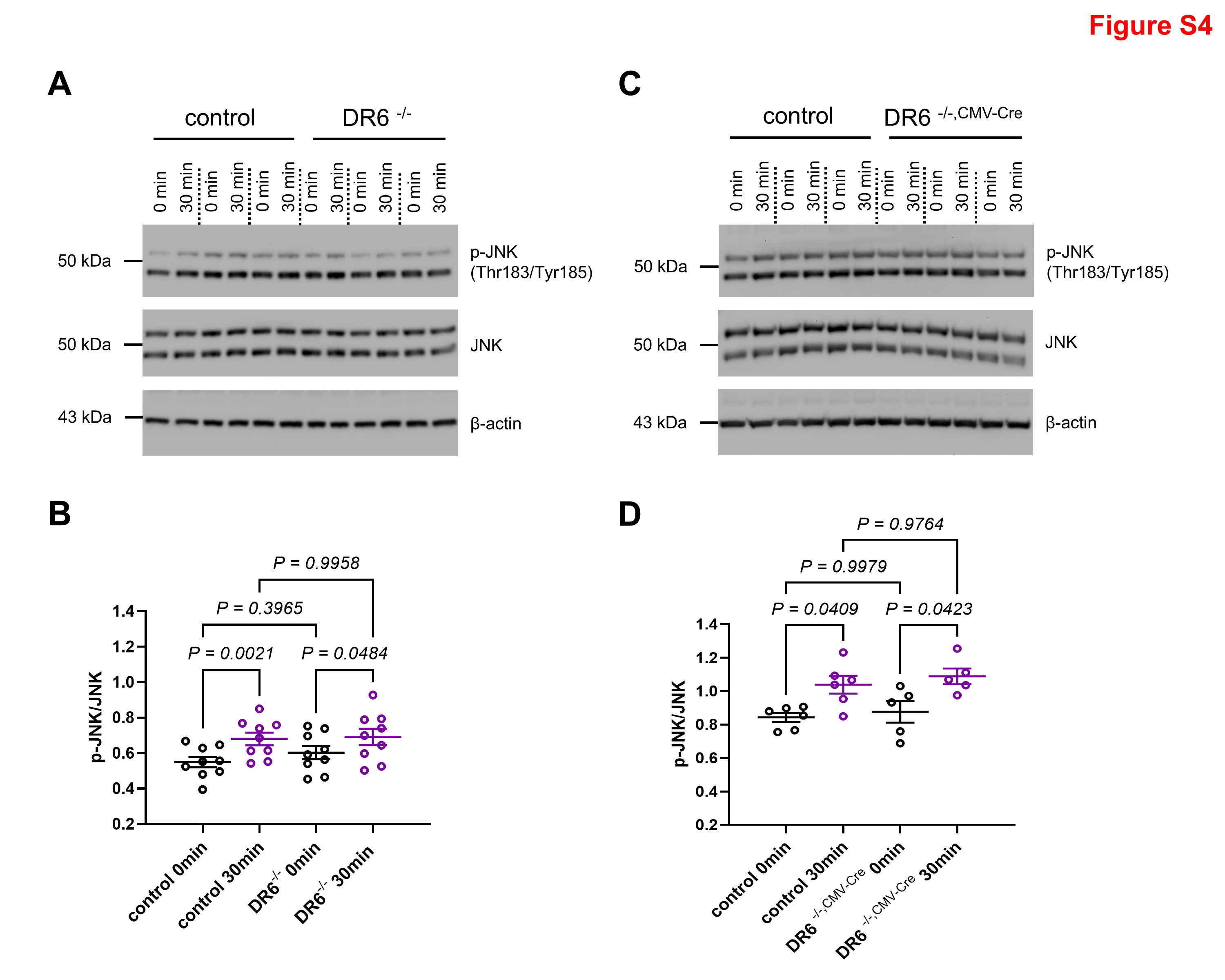

### Figure S5

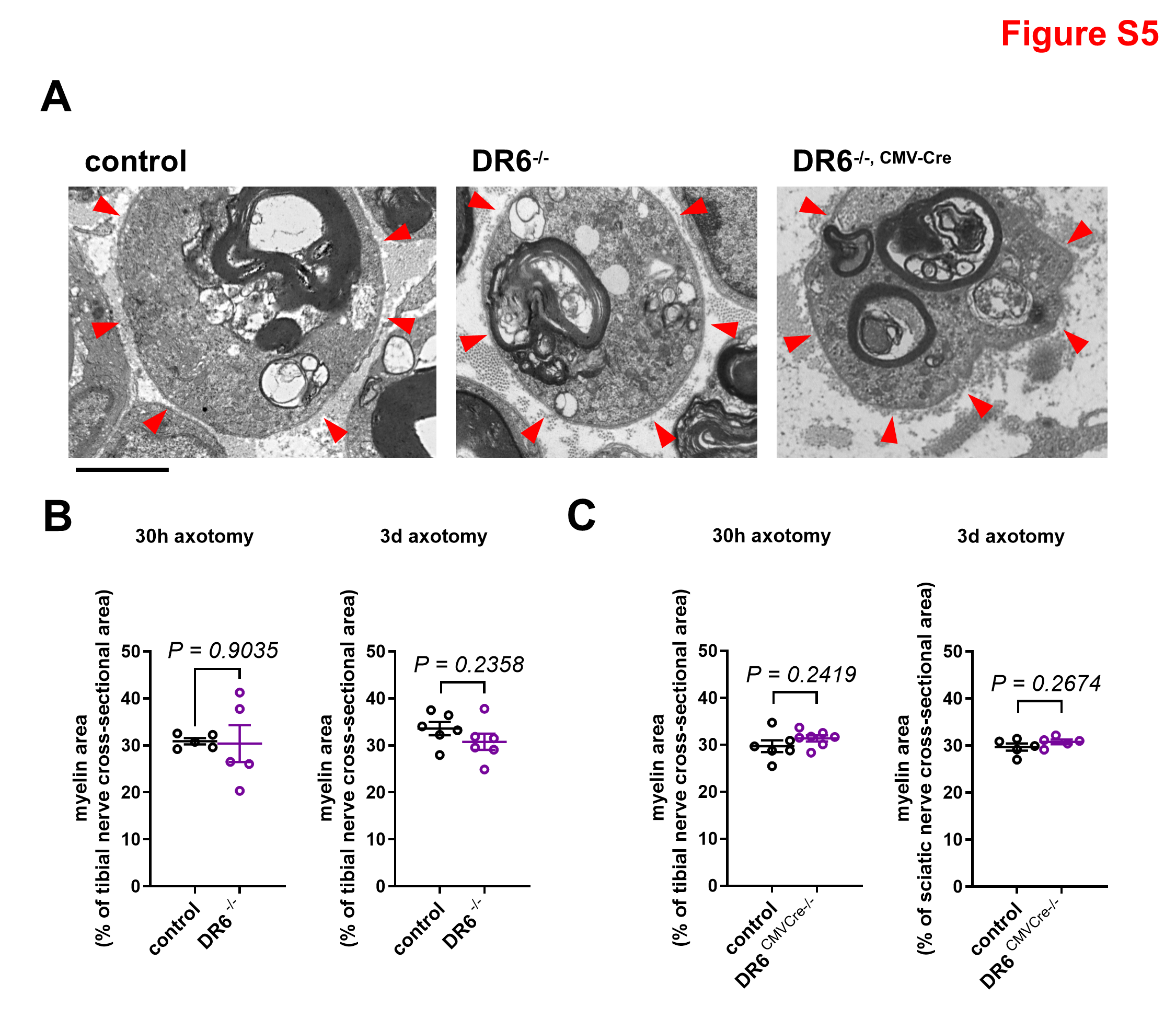
